# Choosing memory retrieval strategies: a critical role for inhibition in the dentate gyrus

**DOI:** 10.1101/2022.05.02.490267

**Authors:** Anne Albrecht, Iris Müller, Aliće Weiglein, Evangelia Pollali, Gürsel Çalışkan, Oliver Stork

## Abstract

Remembering the location of food is essential for survival. Rodents and humans employ mainly hippocampus-dependent spatial strategies, but when being stressed they shift to striatum-mediated stimulus-based strategies. To investigate underlying brain circuits, we tested mice with a heightened stress susceptibility due to a lack of the GABA-synthetizing enzyme GAD65 (GAD65-/- mice) in a dual solution task. Here, GAD65-/- mice preferred to locate a food reward in an open field via a proximal cue, while their wildtype littermates preferred a spatial strategy. The analysis of cFos co-activation across brain regions and of stress-induced mRNA expression changes of GAD65 pointed towards the hippocampal dorsal dentate gyrus (dDG) as a central structure for mediating stress effects on strategy choices via GAD65. Reducing the GAD65 expression locally in the dDG by a shRNA mediated knock down was sufficient to replicate the phenotype of the global GAD65 knock out and to increase dDG excitability. Using DREADD vectors to specifically interfere with dDG circuit activity during dual solution retrieval but not learning confirmed that the dDG modulates strategy choices and that a balanced excitability of this structure is necessary to establish spatial strategy preference. These data highlight the dDG as a critical hub for choosing between spatial and non-spatial foraging strategies.

## 1. Introduction

Remembering the location of a food source is indispensable for survival. Navigating to such a location can be achieved via different neuronal systems dependent on the underlying information processing and operations performed. Rodents and humans deploy two major strategies: They can form an internal map by learning spatial relationships between landmarks or they use a stimulus response strategy, where a sequence of motor events needs to be performed upon specific cues, e.g. turn right at the gas station and then left at the bakery (Packard and Goodman, 2012). While the spatial strategy involves the hippocampus and medial temporal lobe structures, the cue response strategy activates the dorsal striatum (Goodman, 2021; Packard and Goodman, 2013). Moreover, studies in human and rodents demonstrate that prefrontal cortical areas are involved in processing stimulus-reward associations and in choosing specific actions upon these stimuli. Together with cingulate cortical areas, they also process cue- and spatial-related information such as the position of landmarks or of the own position in space. By forming a network with the hippocampus and striatum, prefrontal and cingulate cortical areas mediate choosing spatial or cue-based response strategies when navigating to a reward (Cowen et al., 2012; Kennerley et al., 2006; Rich and Shapiro, 2009; Rudebeck and Izquierdo, 2022).

While a spatial strategy allows for creating shortcuts and is more flexible, individuals under stress favor a cue response learning strategy. To investigate this shift in strategy preferences Schwabe et al. (Schwabe et al., 2008) developed a paradigm where mice had to learn the location of an escape hole on a hole board by either using distal spatial cues for navigation or respond to a bottle located next to the escape hole. While control mice relied on a spatial reward-location strategy, chronically stressed mice preferred responding to the proximal cue. A similar dual solution setup in humans using a computer-based memory game demonstrated that participants with a higher self-reported stress score shifted towards a stimulus-response strategy as well. In a follow-up study, the authors could demonstrate that acute stress, the administration of the stress hormone corticosterone (CORT) and the activation of mineralocorticoid receptors (MR) induce a similar switch of memory systems (Schwabe et al., 2010a, 2010b). However, stress and stress hormones have detrimental effects on hippocampus-dependent learning and long-term potentiation (Roozendaal et al., 2003; Sandi and Pinelo-Nava, 2007) and may already affect spatial strategy preference by impairing spatial learning but not strategy choice during retrieval. Moreover, detailed knowledge about possible molecular mechanisms of stress-induced shifts and their location within the strategy choice neuronal network remains elusive to date.

In our current study we investigated the involved neuronal networks and their potential molecular regulation, focusing on the role of GABAergic interneurons and the key enzyme in GABA synthesis, glutamic acid decarboxylase (GAD). Within the cortex, hippocampus and striatum neuronal activity is shaped by local inhibitory interneurons that use GABA as a neurotransmitter. The GABA-synthetizing enzyme GAD65 critically determines the activity-dependent GABA pools and expression levels of this enzyme are modulated by stress exposure as well as corticosterone (Albrecht et al., 2021b; Bergado-Acosta et al., 2008; Bowers et al., 1998). Mice with a total knock out of GAD65 (GAD65-/-) show increased anxiety, avoidance and hyperarousal as well as a generalized fear memory and deficits in fear extinction (Müller et al., 2015). They provide an interesting model to further elucidate molecular mechanisms and a potential contribution of the inhibitory system to strategy choice in a dual solution paradigm. We therefore compared strategy preferences of GAD65-/- mice and their wildtype littermates in a dual solution task based on locating a food reward in the open field. GAD65-/- mice did not show a spatial strategy preference, even though spatial learning was intact in these animals. Analysis of neuronal co-activation of brain regions via cFos and the analysis of stress-induced expression changes of GAD65 pointed towards the dorsal dentate gyrus (dDG) as a potential hub for mediating strategy choices via GAD65. Accordingly, the local knock down of GAD65 in the dDG reproduced a phenotype reminiscent of the total knock out in the dual solution task and induced an increased excitability of the DG circuit. Using chemogenetic manipulation of the dDG network to dissect effects on learning vs. retrieval confirmed the importance of DG excitation/ inhibition balance for strategy choices during retrieval.

## 2. Material and methods

### 2.1 Animals

All mice were bred and raised in the animal facility at the Institute of Biology, Otto-von-Guericke University Magdeburg. The animals were kept in groups of 2–6 on an inverse 12 h light/dark cycle (lights on at 7 pm with a 30 min dawn phase) with food and water *ad libitum*. Animal housing and animal experiments were in accordance with European regulations for animal experiments and approved by the Landesverwaltungsamt Saxony-Anhalt (Permission Nr. 42502-2-1177 and −1517). GAD65 knock out mice (GAD65-/-) and their wildtype littermates (GAD65+/+) were obtained from a heterozygous breeding and genotyped with allele-specific polymerase chain reaction as described previously (Stork et al., 2000). Further mouse lines engaged were wild type C57BL/6BomTac (M&B Taconic, Germany), SST-Cre (B6(Cg)-Ssttm2.1(cre)Zjh) and PV-Cre (B6;129P2-Pvalbtm1(cre)Arbr/J) mice. Founders of Cre lines were obtained from Jackson laboratories and maintained in a homozygous breeding for SST-Cre and a heterozygous breeding for PV-Cre mice with genotypes being determined with allele-specific polymerase chain reaction using the supplier’s recommendations. From all lines young adult male mice, aged 10-16 weeks, were used for experiments. All experiments were performed during the dark, active phase of the animals and handling was done under red light illumination.

### 2.2 Stress protocol

Animals were randomly assigned to restrained stress or control group. Restrained animals were placed in a plastic cylinder (3 cm diameter x 11.5 cm length) with holes for breathing at the top for 10 min that prevented side movement and restricted movements forth and back. Home cage controls were handled in a fresh cage in parallel. Afterwards all mice were returned to their home cage for 24 h until preparation of trunk blood and brain tissue after decapitation under brief inhalation anesthesia with isoflurane.

### 2.3 Corticosterone ELISA

Trunk blood was collected and allowed to clot at room temperature for 30 min. Clotted material was removed by centrifugation at 3000 rpm for 15 min. Serum was removed immediately and stored at −80°C until further use. Corticosterone serum levels were measured with the IBL Corticosterone ELISA kit (IBL International, Hamburg, Germany) according to the manufacturer’s instructions.

### 2.4 Dual solution task

Five days before the training of a reward location task, animals were food restricted until 85% of the starting body weight was reached. Everyday each mouse received 4 choco krispies (Kellogg Deutschland, Hamburg, Germany) in a plastic lab dish (8.5 cm diameter) within its home cage. For habituation to the training environment, mice were placed for 20 min in an open field (50 cm x 50 cm, 10 lux low light illumination) equipped with patterned paper cuts as distal cues at the wall. After approximately 45 min break in the homecage, mice were reintroduced to the open field, this time equipped with 4 plastic lab dishes (8.5 cm diameter) in the center of each quadrant. One of the dishes contained a wooden toy block (2 cm x 2 cm and 5 cm high) as a proximal cue and a choco krispie placed right next to it. The other 3 dishes contained 1 choco krispie, but access was restricted by a lid with holes for olfactory information. The position of the reward and the proximal cue was randomized across batches but kept consistent over training trials. In each training trial, a mouse was placed randomly in one of the 2 corners equidistant from the dish with the krispie for start and were allowed to explore the arena freely for 4 min. The latency to eat the choco krispie for each trial was assessed as an indicator for learning progress. Each mouse went through 6 training trials with intertrial intervals of approx. 20 min in the home cage.

Animals performed a retrieval trial 24 h after training for which the proximal cue was placed diagonally opposite of the dish which contained the reward previously. Choco krispies were absent in all dishes during retrieval. Again, the respective mouse started the trial in a corner and the first head contact with a dish in a 4 min session was noted by an experimenter blind to genotype and manipulation: a “spatial” choice was made when the mouse entered the previously rewarded dish, a “cued” choice was noted when the dish with the proximal cue was chosen and when any of the other 2 dishes was entered, the response was “neither”. If a mouse did not visit any dish within 4 min, it was classified as “no response”.

Spatial learning abilities of GAD65-/- mice were tested in separate groups, first by applying training and testing without any proximal cue and, second, by presenting a proximal cue during training but not retrieval. As a measure for spatial memory the time spent at the target position during retrieval was used in these modified paradigms.

### 2.5 cFos immunohistochemistry

A subgroup of GAD65-/- mice and their GAD65+/+ littermates received transcardial perfusion with phosphate buffered 4% paraformaldehyde solution at pH 6.8, as previously described (Raza et al., 2017). After postfixation overnight in the same fixative and overnight immersion in wt/vol 20% sucrose at 4°C, brains were cut into 30 μm thick, serial coronar sections at the level of the PFC, posterior ACC/ dorsal striatum and dorsal hippocampus/ amygdala, stored as free-floating sections in phosphate buffered saline at 4°C and then incubated with a rabbit antibody against cFos (1:1000, Cell signaling #2250, Danvers, MA, USA) for 48 h. As secondary antibody biotinylated goat anti-rabbit (1:200, Jackson ImmunoResearch, Ely, UK) in phosphate buffer (PB) with 0.2% Triton (PBT) was used and labelled with Cy2 via Streptavidin (1:1000, Jackson ImmunoResearch, Ely, UK) in PB. Nuclei were visualized with DAPI (300 nM, Thermofischer, Waltham, MA, USA). Immunostainings were imaged using a DMI6000 epifluorescence microscope (Leica, Wetzlar, Germany) in both hemispheres from 2 slices per animal and region. cFos-positive cells were counted manually within the selected brain areas and cell density was normalized to area size measured with imageJ software. The cell counts per target area were averaged for each individual animal and used for statistical comparison between groups.

### 2.6 qPCR

GAD65-/- and GAD65+/+ mice were sacrificed 24 h after restraint stress by decapitation under deep anesthesia with isoflurane. Brains were quickly removed and snap frozen in methylbutane cooled by liquid nitrogen and stored at −80°C. Brains were mounted at a cryostat at −20°C and coronar sections were removed until the start of the dorsal striatum (1.34 mm from Bregma). Striatal tissue was harvested with a stainless-steel sample corer (diameter 0.5 mm, Fine Science Tools, Heidelberg, Germany). Then further sections were removed until the dorsal pole of the hippocampus emerged (starting at −1.34 mm from Bregma) and dDG tissue was harvested with a 0.35 mm diameter sample corer (Fine Science Tools, Heidelberg, Germany). Lysis of tissue samples and isolation of total RNA was performed with the RNeasy Micro Plus kit (Qiagen, Hilden, Germany) according to manufacturer’s instructions. For first-strand cDNA synthesis the Takara PrimeScript RT-PCR Kit (Takara Bio, Shiga, Japan) was used with 50 μM Oligo (dT) and 50 μM random hexamer first strand primers according to the manufacturer’s instructions.

Real-time PCR was performed with a 1:5 dilution of cDNA using QuantStudio3 real-time PCR apparatus (Life Technologies) and TaqMan^®^ reagents with predesigned assays for the target genes GAD65 (alias GAD2, assay ID: Mm00484623_m1), GAD67 (alias GAD1, assay ID: Mm0072566l_sl), MR (alias NR3C2, assay ID: Mm0124l596_ml) and glucocorticoid receptors (GR, alias NR3C1, assay ID: Mm00433832_m1) as well as for the housekeeping gene Glycerinaldehyd-3-phosphat-Dehydrogenase (GAPDH; endogenous control; Life Technologies) that was labeled with another fluorescent dye, allowing for quantitative evaluation. Multiplex PCR samples were run in triplicates with 50 cycles of 15 s at 95°C and 1 min at 60°C, preceded by a 2 min 50°C decontamination step with Uracil-N-glycosidase and initial denaturation at 95°C for 10 min. Gene expression relative to control groups (relative quantification, RQ) was calculated using the ddCT method by normalizing the mean cycle threshold (CT) of each triplicate assay first to the overall content of cDNA using GAPDH as an internal control (dCT; dCT[target gene]) = (CT [target gene]) - (CT [GAPDH]) and then to the control group with ddCT = dCT(sample) – mean dCT (control group). RQ values were obtained by calculating RQ% = (2-ddCT) × 100.

### 2.7 Viral injections

Mice under inhalation anesthesia with isoflurane (1.5-2.5 Vol% in O2) were placed on a stereotaxic frame (World Precision Instruments, Berlin, Germany). 33G injection needles attached to 10 μl NanoFil microsyringes (World Precision Instruments, Berlin, Germany) were placed bilaterally in drillholes at −1.94 mm anterioposterior, ±1.3 mm mediolateral from Bregma and were lowered −1.7 mm dorsoventral from the brain surface into the dorsal dentate gyrus. Viral vectors were injected bilaterally at a volume of 1 μl/side at a flow rate of 0.1 μl/min using a digital microsyringe pump (World Precision Instruments, Berlin, Germany).

To silence GAD65, a small hairpin RNA construct was generated against the oligonucleotide sequence 5⍰-GCATGCTTCCTACCTCTTTCA-3’ (corresponding to NM 008078, base pairs 1599–1619 in mouse and NM_012563.1, base pairs 1337–1357 in rat) and for control a random sequence shRNA (5⍰-TCGTCATGACGTGCATAGG −3⍰) were generated and cloned for production of lentiviral particles as described and validated in Tripathi et al. (2021). Lentiviral particles were injected with a concentration of 10^9^ particles/μl. Virus spread was determined by the expression of the marker GFP (see Suppl. Fig. 1). The knock down efficiency for GAD65 in the mouse hippocampus was determined by western blot protein expression (see Suppl. Fig. S2).

For chemogenetic manipulation of DG circuits designer receptors exclusively activated by designer drugs (DREADDs)-based tools were employed. The following DREADD adeno-associated viruses (AAV), which were gifts from Bryan Roth and obtained via Addgene (Watertown, MA, USA), were used: pAAV-CaMKIIa-hM4D(Gi)-mCherry (Addgene viral prep #50477-AAV8; http://n2t.net/addgene:50477); RRID:Addgene_50477) and pAAV-CaMKIIa-GFP (Addgene viral prep #50469-AAV8; http://n2t.net/addgene:50469; RRID:Addgene_50469) were used for silencing excitatory cells in the DG. hSyn-DIO-hM4Di-mCherry (Addgene viral prep #44362; http://n2t.net/addgene:44362; RRID:Addgene_44362), AAV hSyn-DIOhM3Dq-mCherry (Addgene viral prep #44361; http://n2t.net/addgene:44361; RRID:Addgene_44361) and AAV hSyn-DIO-mCherry (Addgene viral prep #50459; http://n2t.net/addgene:50459; RRID:Addgene_50459) were used for chemogenetically manipulating interneurons in the respective cre-transgenic lines (Raza et al., 2017). AAVs were injected at > 5×10^12^ particles/μl.

For activation of chemogenetic hM4Di and hM3Dq receptors, animals received an i.p. injection of 10 mg kg-1 clozapine-N-oxide (Enzo Life Sciences, Germany) in physiological saline 1 h before either retrieval or training of the dual solution task.

For histological verification of viral vector expression, animals were transcardially perfused at the end of the experiment and 3-4 alternating 30 μm sections of the dorsal hippocampus (from Bregma AP: −1.58 to AP: −2.18 mm) were imaged using a DMI6000 epifluorescence microscope (Leica, Wetzlar, Germany). As observed previously, marker expression is expected in the dendrites and axons of dDG granule cell (Raza et al., 2017). Only mice with a correct bilateral marker expression in the molecular layer and the mossy fibers of the dDG were therefore considered for behavioral experiment.

### 2.8 Slice electrophysiology

14 to 16 days after the shRNA-mediated viral knock down of GAD65 in the dorsal DG, mice were decapitated under deep anesthesia with isoflurane and their brains were quickly removed and submerged into cold (~4°C) carbogenated (5% CO_2_/95% O_2_) artificial cerebrospinal fluid (aCSF, composition in mM: 129 NaCl, 21 NaHCO_3_, 3 KCl, 1.6 CaCl_2_, 1.8 MgSO_4_, 1.25 NaH_2_PO_4_, 10 glucose). Parasagittal brain slices including the transverse-like dorsal hippocampal sections (~400 μm) were cut from the dorsal pole on an angled platform of approx. 12°. Three to four slices per hemisphere were transferred to an interface chamber perfused with aCSF at 30 ± 0.5°C (flow rate: 1.8 ± 0.2 mL/min, pH 7.4, osmolarity ca. 300 mOsmol/kg) and incubated at least for 90 min before extracellular field recordings were performed.

For dDG electrophysiology (Fig. 3Ci), a bipolar tungsten wire stimulation electrode (with exposed tips of approx. 20 μm, tip separations of approx. 75 μm, electrode resistance in ACSF: approx. 100 kΩ) was placed at the perforant path (PP) presumably stimulating both medial and lateral fibers. Population spike (PS) and positive-going field postsynaptic potential (fEPSP) were recorded with a borosilicate glass electrode filled with aCSF (1-2 MΩ) from granule cell layer of DG at 70–100 mm depth as previously described (Albrecht et al., 2016). Stable PP-induced responses were verified for ~10 min (0.033 Hz with pulse durations of 100 μs) before obtaining an input–output (I-O) curve (Simulation intensities in μM: 10, 20, 30, 40, 50, 75, 100, 150, 200). Paired pulse (PP) responses (Stimulus intervals in ms: 10, 25, 50, 100, 250, 500) were recorded using the ~50-60 % of the maximum PS amplitude.

Signals were sampled at a frequency of 5 kHz, pre-amplified with a custom-made amplifier, low-pass filtered at 3 kHz and stored on a computer hard disc for offline analysis. Evoked potentials PS and fEPSP were analyzed using MATLAB (MathWorks, Natick, MA) and Spike2-based analysis tools (CED, Cambridge). For the PP-DG electrophysiology, the slope of positive-going fPSP was analyzed by calculating the slope (V/s) between the 20 and 80% of the fEPSP amplitudes. The DG PS amplitude was measured by calculating the average of peak-to-peak amplitude (mV) of descending and ascending phase of PS. For calculating the paired-pulse ratio, the magnitude of the second evoked potential was divided by the first one. For methodological details on mossy fiber-to-CA3 synapse or Schaffer collateral-to-CA1 synapse electrophysiology see supplementary material (Suppl. Fig. 3 and 4).

### 2.9 Statistics

For all experiments, normal distribution of the data was assessed with Shapiro-Wilk’s test. Equality of variances was tested using Levene’s test. Comparisons of two groups were made with Student’s t-test or with Mann-Whitney U-test, in case of non-normal distribution of the data. In case of unequal variances, a Welch test was applied. For comparing measures over time (e.g. during dual solution learning), stimulation intensities or intervals (during slice recordings), repeated measure ANOVA was applied, if necessary with Greenhouse-Geißer corrections for sphericity issues. Correlation of cFos intensities between areas were assessed using Pearson’s correlation coefficient. For comparing the distribution of strategy choices between groups the likelihood-ratio chi-square (X^2^) test, optimized for small sample sizes, was used. For statistical analysis of the data GraphPad Prism software (version 9, SD, California) and SPSS (version 28, IBM, New York) was used. Differences were considered statistically significant if p-value (p) < 0.05.

## 3. Results

### 3.1 GAD65-/- mice preferred a cue-based strategy in the dual solution reward memory task

To assess the stress susceptibility of GAD65-/- mice a brief submission to 10 min restrained stress was performed and CORT serum levels were determined 24 h in trunk blood (GAD65+/+ control: n=7; GAD65+/+ restrained: n=6; GAD65-/- control: n=6; GAD65-/- retrained: n=5). CORT levels were elevated in GAD65-/- irrespective of stress (Fig. 1A; Mann-Whitney-U-test for main effects in non-normal distributed data set: U=106, p=0.047). Paired comparisons revealed that GAD65-/- mice had higher serum levels of CORT than their wildtype littermates under control conditions (control GAD65+/+ vs. -/-: U=38, p=0.014), while 24 h after restrained stress no genotype differences were observed (restrained GAD65+/+ vs. -/-: U=18, p=0.662).

**Fig. 1:**
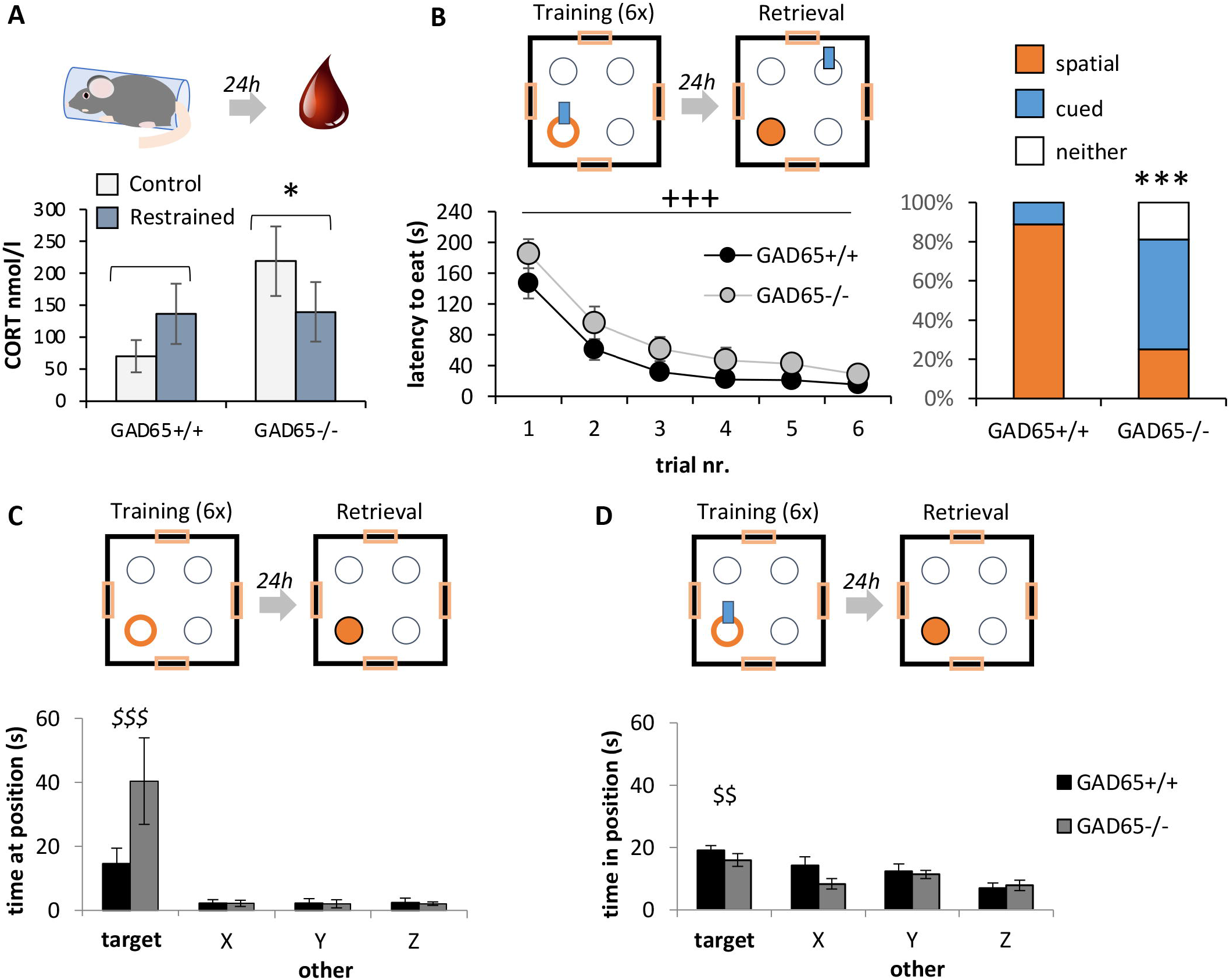
Strategy preference in GAD65-/- mice. (**A**)GAD65-/- mice showed elevated corticosterone serum levels already under baseline control conditions that were not further elevated 24 h after a brief restraint stress episode, suggesting altered stress susceptibility. (**B**)In the dual solution task (scheme on top), GAD65-/- mice showed a good learning performance of the reward location over 6 trials. When tested for their strategy preference 24 h later, they did not display a spatial preference in comparison to their wildtype littermates. (C) When the experimental setup only allowed for a spatial solution in order to learn the reward location, GAD65-/- mice showed a spatial preference for the reward location during retrieval. (D) Similarly, when a cue was present during training but not during retrieval, GAD65-/- mice still displayed a spatial preference similar to their wildtype littermates. All values means ± sem. * significant difference between genotypes, p<0.05; *** p<0.001; +++ significant learning effect over trials, p<0.001; $$ significant difference of target position compared to other positions, p<0.01; $$$ p<0.001.

Both, GAD65-/- (n=18) and GAD65+/+ (n=16) mice learned the food location well during training in the dual solution task, with a trend to lower latencies in wildtype mice (Fig. 1 B; Repeated Measure ANOVA with Greenhouse-Geißer correction: trials F(2.466, 78.913)=56.333, p<0.001; trials x genotype F(2.466, 78.913)=0.405, p=0.711; genotype F(1,32)=3.831, p=0.059). During retrieval, GAD65+/+ mice showed a preference for a spatial response strategy, while GAD65-/- showed a more diverse response pattern (Fig. 1B; likelihood-ratio chi square test X^2^(2)=16.569, p=0.001).

When no proximal cue was present during training and retrieval, mice of both genotypes (GAD65+/+: n=9; GAD65-/-: n=7) spent most of the time during the retrieval in the target position previously probed with the reward (Fig. 1C; Repeated Measure ANOVA with Greenhouse-Geißer correction for position: F(1.067, 14.942)=15.358, p=0.001; with p<0.01 for target vs. other positions, LSD posthoc test). No main effects of genotype (F(1,14)=2.949, p=0.108) nor significant interactions of position x genotype: F(1.067, 14.942)=4.062, p=0.06) were observed. GAD65-/- mice showed no significantly increased time at the target position compared to their wildtype littermates (paired comparison, Mann-Whitney-U-test: U=44.5, p=0.174). When a cue was present during training but not retrieval, mice of both genotypes (GAD65+/+: n=8; GAD65-/-: n=10) spent more time in the target position (Fig. 1D; Repeated Measure ANOVA for position: F(3, 66)=12.255, p<0.001; with p<0.01 for target vs. other positions, LSD posthoc test) and no effects of genotype (F(1,22)=1.555, p=0.225) or interactions of position x genotype: F(3,66)=1.463, p=0.233) were observed. Thus, while stress-susceptible GAD65-/- mice lost the spatial preference observed in GAD65+/+ mice in a dual solution task, they were able to acquire spatial information for the reward location and could use this information for retrieval if no other option was available.

### 3.2 Interregional interaction and GAD65 expression regulation after stress point towards the dDG as a hub for stress-related effects

After the retrieval, the number of cFos-positive neurons (per area) was counted in selected brain regions of spatial-preferring GAD65+/+ mice (n=5) vs. GAD65-/- with a cue-based retrieval choice (n=5). No significant differences were observed in any of the selected areas (Fig. 2A; T-/ Mann-Whitney-U-Tests for PL/IL: T(8)=0.843, p=0.424; ACC: T(8)=-0.141, p=0.891; striatum: T(8)=-0.211, p=0.838; DG: T(8)=0.583, p=0.576; CA1: T(8)=0.318, p=0.758; CA3: T(8)=-0.539, p=0.604; LA: T(8)=0.912, p=0.388; BLA: T(8)=0.458, p=0.659; CeA: U=14, p=0.814). However, a co-activation analysis using Pearson’s correlation co-efficient revealed negative correlations between the prefrontal areas (PL/IL) or the ACC and the dDG, as well as a positive correlation between the dDG and the central amygdala in GAD65+/+ mice with a spatial preference. In GAD65-/- mice the correlation patterns of the DG with prefrontal cortical areas, the striatum and the amygdala appeared reversed between groups (see Fig. 2B for color-coded correlation coefficients).

**Fig. 2:**
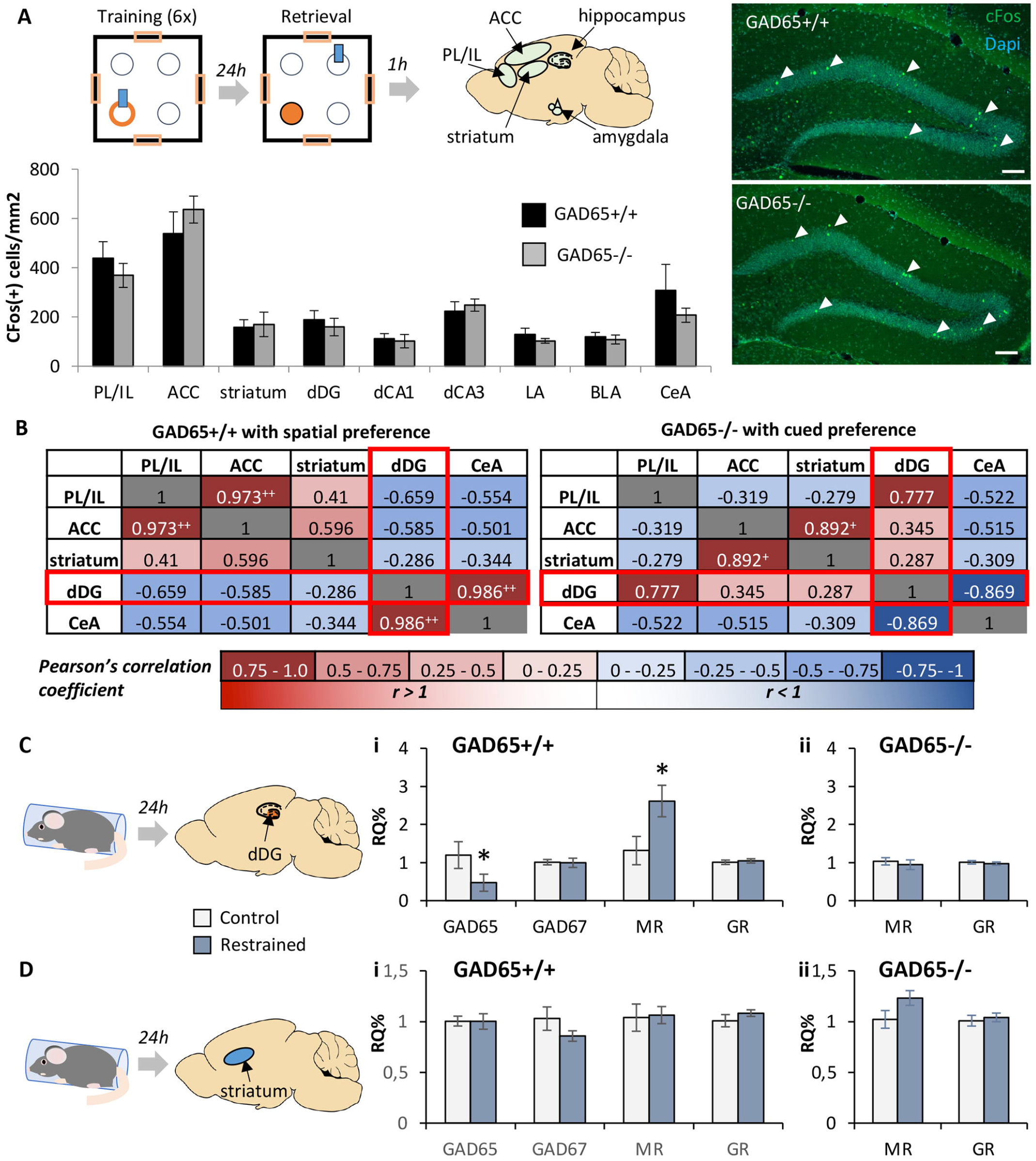
Interregional activation patterns during dual solution retrieval and GAD65 expression regulation after stress point towards the dDG as a hub for stress-related effects. (**A**)GAD65-/- mice and their wildtype littermates were trained in the dual solution task and perfused 1 h after retrieval brains for subsequent cFos analysis. Example images are shown for cFos labeling (green: cFos; cyan: DAPI nuclear staining; white arrow head: exemplary cFos-positive cell; scale bar = 100 μm) in the dorsal dentate gyrus. No difference in the number of cFos positive cells was evident in any of the regions analyzed between GAD65 +/+ mice preferring a spatial solution (n=5) and GAD65-/- mice using cue-based strategy (n=5). (B) Interregional correlations of cFos activation after DS retrieval and strategy choice demonstrated shifts in co-activation of the dDG together with PL/IL, and CeA in between the groups. (C) The dDG was also susceptible to gene expression changes induced by restrained stress. In (i) GAD65+/+ mice GAD65 mRNA levels were downregulated and mineralocorticoid receptor (MR) upregulated 24h after brief restrained stress, while (ii) MR expression was unaltered in GAD65-/- mice (n=5-7 per group). (**D**)In comparison, no stress-induced expression changes were observed in the striatum of (i) GAD65+/+ and (ii) GAD65-/- mice (n=5-7 per group). Expression of the isoform GAD67 and of the glucocorticoid receptor (GR) remained unchanged in both regions. All values means±sem. + significant Pearson’s correlation, p<0.05; ++ p<0.01; * significant difference between genotypes, p<0.05.

**Fig. 3:**
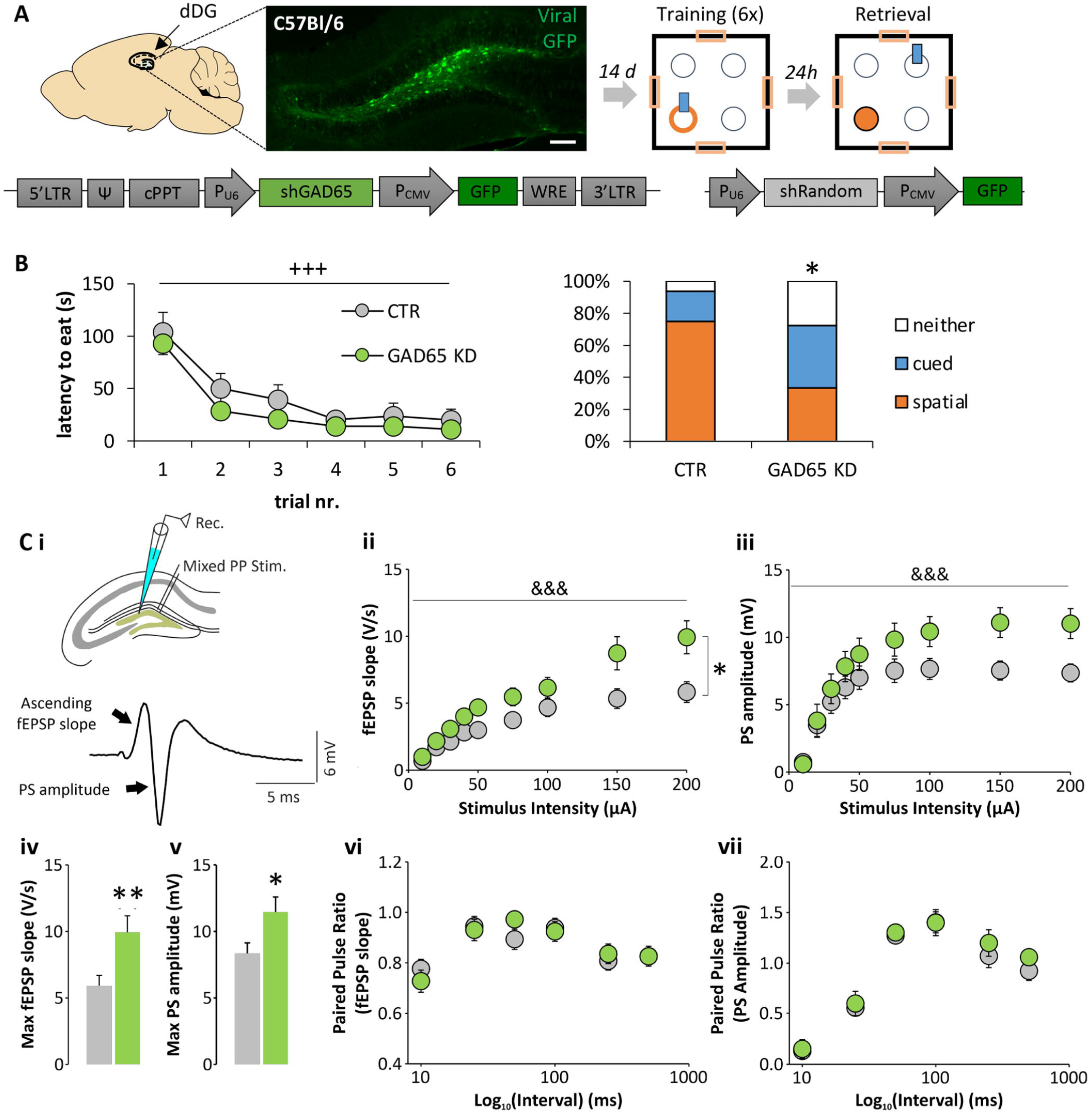
Local knock down of GAD65 in the dDG leads to loss of spatial preference in a dual solution task. (**A**)Vector scheme and workflow for local knock down of GAD65 in the dDG before dual solution training and retrieval. (B) The local knock down of GAD65 in the dDG is sufficient to replicate the phenotype of total GAD65-/- mice. (C) Electrophysiological profile of the dDG after local GAD65 knock down (CTR: n=18 slices from 9 animals; GAD65 KD: n=12 slices from 8 animals), (i) Scheme for perforant-path induced somatic field postsynaptic potential (fEPSP) and population spike (PS) responses. Input-output (I/O) curves demonstrate an enhanced excitability after GAD65 KD in the dDG evident by (ii) increased fEPSP slope and (iii) PS amplitudes, (iv) Accordingly, the maximum evoked fEPSP slope and (v) PS amplitude is increased in GAD65 KD slices. No differences in the paired pulse ratios were observed for (vi) the fEPSP slopes or (vii) the PS amplitudes. All values means±sem. +++ significant learning effect over trials, p<0.001; * significant difference from CTR group, p<0.05, ** p<0.01, *** p<0.001. &&& p<0.001, significant stimulus intensity effect.

An important role of the dDG in mediating stress-related effects on dual solution strategy choice was further supported by the expression analysis 24 h after restrained stress (n=5-6 for control, n=6 for restrained). Such a brief stressor before training has been shown to shift retrieval strategies, at least in part via CORT signaling on mineralocorticoid receptors (MR). 24 h after restrained stress, mRNA levels of MR were increased in the dDG of GAD65+/+ mice (Fig. 2Ci; T(10)=-2.343, p=0.041), while the other receptor for CORT, GR, remained unchanged (T(10)=-0.433, p=0.674). Moreover, in wildtype mice restrained stress lastingly reduced the expression of GAD65 in the dDG (Fig. 2Ci; Mann-Whitney-U-test, U=3, p=0.03), but not GAD67 (T-test, T(9)=0.114, p=0.912). The expression of MR and GR was not altered in the dDG of GAD65-/- mice 24 h after restrained stress (Fig. 2Cii; MR: T-test T(11)=0.572, p=0.579; GR: T-test T(11)=0.541, p=0.6). In the dorsal striatum no changes in expression levels were observed in GAD65+/+ mice (Fig. 2Di; GAD65: Welch-tests for unequal variances, T(8.212)=0.041, p=0.968; GAD67: T-test T(10)=1.361, p=0.203; MR: T(10)=-0.157, p=0.878; GR: T(10)=−1.09, p=0.301) and in GAD65-/- mice (Fig. 2Dii; MR: T-test T(11)=−1.817, p=0.097; GR: T-test T(11)=-0.466, p=0.65).

Together, these data suggested a relevant role for the dDG in stress responses and GAD65 expression regulation in wildtype mice that prompted us to specifically target this region in the dual solution task.

### 3.3 Local knock down of GAD65 in the dDG is sufficient to induce a loss of spatial strategy preference in the dual solution task

Based on the co-activation analysis, we then used a recently established shRNA-mediated lentiviral vector for GAD65 knock down and injected it into the dDG (Tripathi et al., 2021). Two weeks after virus injection training in the dual solution task took place. Mice with the local GAD65 knock down (GAD65 KD, n=18) acquired the task as successfully as mice injected with a control vector containing a random sh sequence (CTR, n=16; Fig. 3B; Repeated Measure ANOVA with Greenhouse-Geißer correction: trials: F(2.023,64.734)=46.477, p<0.001; trials x group: F(2.023,64.734)=0.426, p=0.657; group: F(1,32)=1.386, p=0.248). Different strategy choice patterns were visible during retrieval 24h later (Fig. 3B; X^2^(2)=6.478, p=0.039), with CTR but not GAD65 KD mice showing a spatial preference.

To assess the impact of GAD65 KD on excitability of the dDG, perforant-path induced fEPSP and PS responses were recorded in the DG granule layer in slices prepared from a different set of CTR and GAD65 KD mice (Fig. 3Ci). Analysis of I/O curves revealed an increased somatic fEPSP slopes in GAD65 KD slices (Fig. 3Cii; Repeated Measure ANOVA with Greenhouse-Geißer correction: stimulation intensity F(1.397,39.12)=86.93, p<0.0001; stimulation intensity x group: F(8,224)=6.352, p<0.0001; group: F(1,28)=5.965, p=0.0212)and PS amplitudes (Fig. 3Ciii; Repeated Measure ANOVA with Greenhouse-Geißer correction: stimulation intensity F(2.396, 67.09)=63.80, p<0.0001; stimulation intensity x group: F(8,224)=3.205, p=0.0018; group: F(1,28)=2.931, p=0.0979). Accordingly, the maximum evoked somatic fEPSP slope (Fig. 3Civ; Mann-Whitney-U-test, U=39, p=0.0027) as well as PS amplitude (Fig. 3Cv; T-test, T(28)=2.327, p=0.0274) were increased in slices from GAD65 KD mice. When determining paired pulse responses as a measure of short term plasticity, no significant differences were observed after GAD65 KD for the somatic fEPSP slopes (Fig. 3Cvi; Repeated Measure ANOVA with Greenhouse-Geißer correction: interval F(3.361, 87.39)=17.12, p<0.0001; interval x KD F(5,130)=1.366, p=0.2411; KD F(1,26)=0.01234, p=0.9124) and for the PS amplitudes (Fig. 3Cvii: Repeated Measure ANOVA with Greenhouse-Geißer correction: interval: F(1.664, 46.59)=64.26, p<0.0001; interval x group: F(5,140)=0.2413, p=0.9435; group: F(1,28)=0.5033, p=0.4839).

Of note, we found no evidence for altered neurotransmission and plasticity in the MF-CA3 (Suppl. Fig. 3) and SC-CA1 (Suppl. Fig. 4) synapses in hippocampal slices of GAD65 KD mice, indicating that GAD65 KD induced changes in the excitability of dDG remain local without altering baseline synaptic transmission and plasticity in the downstream regions of the hippocampal trisynaptic circuit.

Taken together, a local GAD65 knock down in the dDG induced a similar phenotype in the dual solution task as the total knock out of GAD65 and increased the excitability of this structure, thereby suggesting an important role of dDG excitability in spatial strategy preference.

### 3.4 Modulation of dDG excitatory activity during retrieval affects strategy choices

To dissect whether inhibition by dDG interneurons modulates strategy choices during dual solution task retrieval, chemogenetic activation and inhibition of interneurons before retrieval were performed via DREADDs in interneuron-specific cre-driver lines (Fig. 4A). SST-Cre mice were used to target somatostatin-positive (SST(+)) interneurons in the dDG and to increase (hM3Dq, n=8) or reduce (hM4Di, n=8) their activity, in comparison to controls (CTR, n=9). The groups did not differ in their learning performance, as expected (Fig. 4B; Repeated Measure ANOVA with Greenhouse-Geißer correction for trial F(2.784,61.243)=18.357, p<0.001; trial x group F(5.568,61.243)=0.846, p=0.533; group F(2,22)=1.934, p=0.168). Modulating SST(+) interneuron activity by CNO ligand injection before retrieval induced no differences and all groups showed spatial dominating response patterns (Fig. 4B, upper panel; X^2^(4)=0.196, p=0.906). Manipulating parvalbumin-positive (PV(+)) interneurons was achieved by injecting activating (hM3Dq, n=10), inactivating (hM4Di, n=10) and control constructs (CTR, n=9) in the dDG of PV-Cre-mice and applying CNO again 1 h before retrieval. As expected, training 24 h earlier was not affected (Fig. 4B, lower panel; Repeated Measure ANOVA with Greenhouse-Geißer correction for trial F(1.788,50.063)=42.462, p<0.001; trial x group F(3.567,50.063)=1.381, p=0.256; group: F(2,28)=1.793, p=0.185). Activating or inactivating PV(+) did not shift response patterns significantly either (Fig. 5C; X^2^(2)=3.714, p=0.446). However, spatial preference was less established in control animals of this strain (50% in PV CTR) compared to all other control mouse cohorts used in this study, suggesting strain effects.

**Fig. 4.**
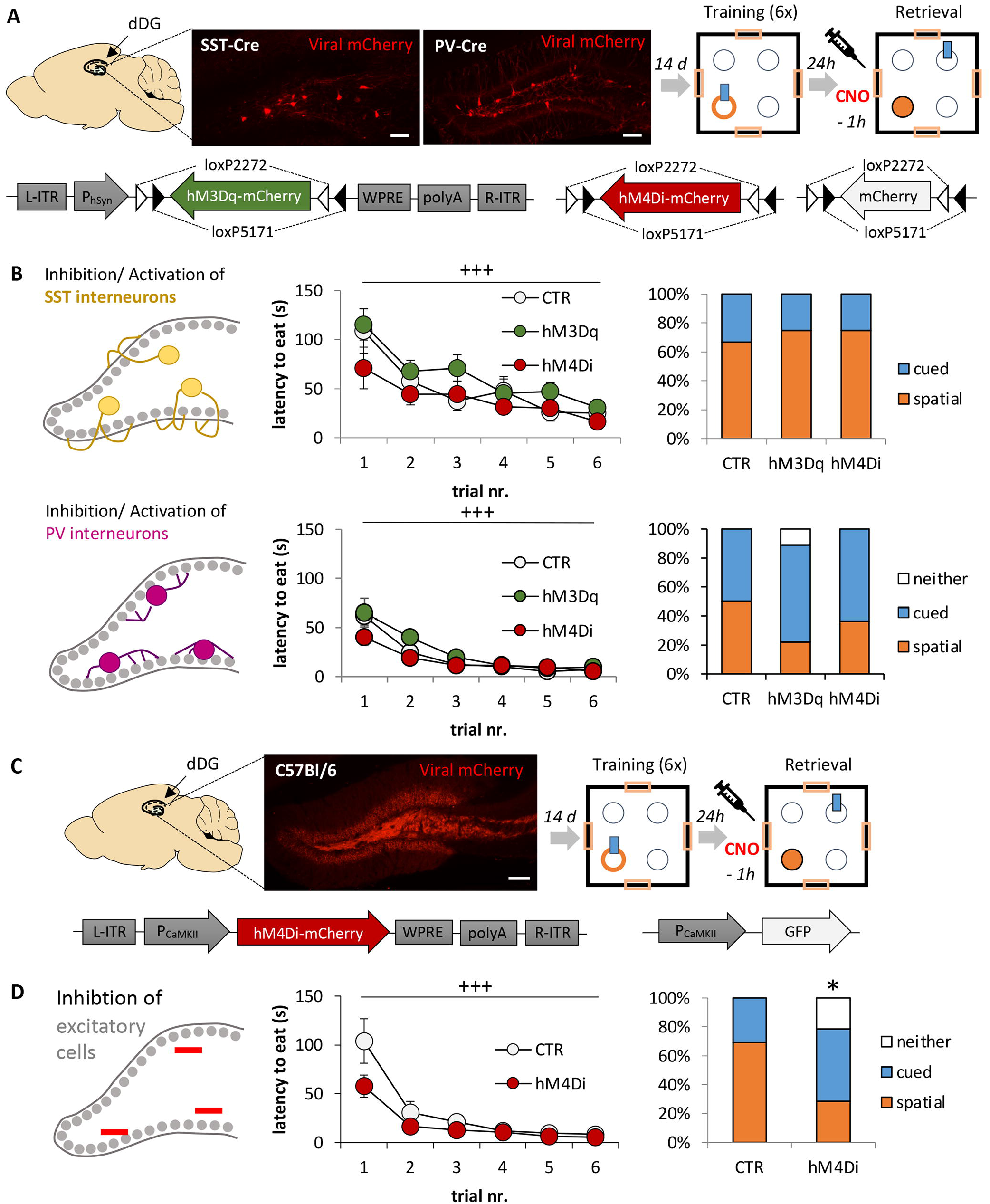
Modulation of dDG excitatory activity during retrieval affects strategy choices. (**A**)Scheme for chemogenetic activation/ inhibition of somatostatin (SST)- and parvalbumin (PV)-positive interneurons during dual solution retrieval. (**B, upper panel**)Activation or inhibition of dDG SST(+) cells during dual solution retrieval did not affect strategy choice (CTR: n=9; 3Dq: n=8; 4Di: n=8). (**B, lower panel**)Activation or inhibition of PV(+) interneurons during dual solution retrieval did not affect strategy choice significantly, but spatial strategy preference appeared diminished in the PV-hM3Dq group (not significant; CTR: n=9; 3Dq: n=10; 4Di: n=10). (C) For comparison, chemogenetic inhibtion of excitatory dDG cells was achieved using CamKII-hM4Di constructs. Expression of the marker mCherry is observed in the molecular layer and mossy fibers, containing dendritic and axonal compartments of dDG granule cells. (**D**)Inhibiting excitatory cell activity during retrieval induced a strategy shift with a loss of spatial preference (n=l4; CTR: n=l3). All groups did not differ in their training performance. All values means±sem. +++ significant learning effect over trials, p<0.001; * significant difference to CTR group, p<0.05.

For comparison, dDG excitatory cells were chemogenetically inhibited before retrieval by using inhibitory DREADD constructs under the CamKII promotor (Fig. 4C; hM4Di, n=14; vs. control construct, n=13). Both, hM4Di and the CTR groups learned the location of the reward well during training (Fig. 4B; Repeated Measure ANOVA with Greenhouse-Geißer correction for trial F(1.476,36.9)=28.779, p<0.001; trial x group interaction F(1.476,36.9)=2.642, p=0.099; group F(1,25)=3.919, p=0.059). After inactivating excitatory DG cells during retrieval, strategy choices were shifted from spatial pattern in CTR to cue-based strategy in the hM4Di groups (Fig. 4D; X^2^(2)=6.924, p=0.031). Noteworthy, inhibiting DG activity before training did not induce a shift in strategy choice (Suppl. Fig. 5). Together, a direct contribution of either SST- or PV-positive DG interneuron populations to strategy choices during dual solution retrieval could not be confirmed, despite their potential role in modulating DG excitability. Targeting, excitatory cells of the dDG during retrieval, however, induced a strategy shift, suggesting a specific role of this region in strategy choice during retrieval rather than in acquiring strategies during training.

## 4. Discussion

The adaptation to stress involves various neurophysiological and behavioral responses, including a switch from previously acquired spatial to cue-based search strategies (Schwabe et al., 2010a, 2010b, 2008). Our findings suggest that these alterations involve a regulation of inhibitory functions in the dDG, via regulation of the GABA synthesizing enzyme GAD65, and excitability of the DG granule cells.

As one of the two isoenzymes synthetizing GABA, GAD65 expression is regulated in an activity-dependent manner and is modulated by stress exposure (Albrecht et al., 2021b; Bergado-Acosta et al., 2008; Bowers et al., 1998). GAD65-/- mice display a behavioral phenotype associated with a maladaptation to challenging environments, including increased anxiety, increased fear memory as well as a heightened generalization of conditioned fear and reduced fear extinction (see (Müller et al., 2015), for review). In the current study increased plasma levels of corticosterone were observed in GAD65-/- mice handling controls, already without previous exposure to restrained stress. While basal differences in corticosterone levels had not been observed in previous studies (Stork et al., 2003), likely due to differences in rearing and analysis conditions including circadian rhythm, they further support the notion of an altered stress response system in these mice.

Stress and elevated CORT levels have been shown to induce a shift in strategy preference in mice favoring stimulus-response based strategies (Schwabe et al., 2010a). We therefore tested unstressed, naïve GAD65-/- mice in a dual solution foraging task, based on locating a food reward in an open field. Like in other protocols based on aversive and appetitive stimuli (see (Goodman, 2021), for review), mice have the option to build either a spatial map and navigate in relation to distal cues like signs at the walls or to respond to cues placed proximal to the goal, in our case the food reward. After establishing both strategies during the memory acquisition phase, the preferred solution strategy can be assessed during retrieval by moving the proximal cue to a different position. Mice will then either direct their search to the cue or to the trained location (spatial solution). GAD65+/+ mice preferred a spatial strategy in that case, as seen in wildtype mice in other dual solution paradigms e.g. using a holeboard (Schwabe et al., 2008). GAD65-/- mice, however, showed a cued response in 50% of the individuals, as observed previously in chronically or acutely pre-stressed mice (Schwabe et al., 2008, 2009). Importantly, GAD65-/- mice showed in our protocol no aversion against novel food nor any alterations in activity or anxiety-like behavior in the open field (see Suppl. Fig. 6, 20-min habituation before the dual solution training), thereby rendering confounding effects of heightened anxiety on learning or retrieval unlikely.

Across different dual solution paradigms, stress hormones such as CORT and activation of the basolateral amygdala have been demonstrated to induce stress-related shifts of strategy choices (Packard, 2009; Packard and Goodman, 2012). As these factors may impair spatial memory acquisition (Roozendaal et al., 2003; Sandi and Pinelo-Nava, 2007), we tested whether GAD65-/- mice have a spatial learning deficit. However, these mice were able to acquire spatial memory for a food reward when only a spatial solution was available or when the proximal cue was absent during the retrieval. This suggests that GAD65-/- mice have no spatial learning or memory deficit per se, in line with undisturbed context processing in aversive fear conditioning paradigms (Bergado-Acosta et al., 2008; Sangha et al., 2009). Together, unstressed GAD65-/- mice prefer a cue-based response strategy just like stressed mice in previous studies, but neither disturbed emotional processing nor spatial learning deficits appear accountable for this effect.

Previous studies suggest that both systems, hippocampus and dorsal striatum, are activated during dual solution training and their cellular activity relates to strategy choices (Kathirvelu and Colombo, 2013; Yagi et al., 2017). Studies in humans and rodents show that decision making processes such as choosing an appropriate strategy also depend on the mPFC and ACC (Rich and Shapiro, 2009; Rudebeck and Izquierdo, 2022). Using the neuronal activity marker cFos, we mapped neuronal activation in these brain regions and found no differences in neuronal activation patterns of spatial-preferring GAD65+/+ vs. cued-preferring GAD65-/- mice, further supporting that hippocampus and striatum-based retrieval systems are intact in GAD65-/- mice. However, when correlating activities between brain areas, we noted a shift in the co-activation specifically of the dDG with prefrontal cortical areas and the central amygdala areas between genotypes, indicative of a change in functional connectivity.

The dDG is perfectly suited to integrate cognitive processes and emotional arousal and plasticity and is modulated by stress and amygdala activation (Albrecht et al., 2021a; Fa et al., 2014). Information processing in the DG is achieved by strong inhibition via GABAergic interneurons that form local networks with the excitatory granule cells (Acsády and Káli, 2007) and may mediate stress effects and the impact of GAD65 deficiency on strategy preferences. We therefore assessed whether a brief restrained stress exposure, a procedure that induced a shift in strategy choice in previous studies (Schwabe et al., 2010b), alters the expression of GAD65 in the dDG of wildtype mice. Indeed, GAD65 mRNA expression was reduced in the dDG while remaining unaltered in the striatum of GAD65+/+ mice. Moreover, an increased expression of MR was observed in the dDG of GAD65+/+ mice, but not in their striatum. In GAD65-/- mice MR levels remained unchanged in the dDG and only a slight, but insignificant increase in MR expression was visible in the striatum, further supporting an altered regulation of the stress axis in GAD65-/- mice. The activation of MR has previously been linked to the stress-induced strategy shift in dual solution paradigms. An orally administered MR antagonist could prevent stress-induced shifts in strategy choices (Schwabe et al., 2010a), while mice overexpressing MR in excitatory neurons showed a stronger switch to cued response strategies after stress (ter Horst et al., 2012). Thus, increased MR levels in the dDG of wildtype mice after stress underline a possible central role of the dDG.

Accordingly, we then commenced by mimicking a molecular aspect of stress exposure by reducing the expression of GAD65 only in the dDG using lentiviral vectors. This treatment clearly replicated the lack of spatial strategy preference observed in the total GAD65-/- mice. Field recordings in DG slices from such animals demonstrated that a GAD65 knock down furthermore increased excitability of dDG circuits without altering paired pulse responses. Previous studies in GAD65-/- mice found a reduced paired pulse inhibition, but only in the CA1 area (Tian et al., 1999). The current results suggest a heightened excitability of the DG, in accordance with a reduced inhibitory activity due to GAD65 knock down.

Two major populations of GABAergic interneurons shape the local circuits of the DG. We next investigated therefore their potential impact on strategy choices in the dual solution task by chemogenetic intervention. This approach also allowed for a targeted manipulation of the training vs. retrieval phase. First, we manipulated somatostatin interneurons, which are predominantly located in the hilus of the dDG and mediate inhibition of distal dendrites of excitatory granule cells (Tallent, 2007). These cells control the size of engrams for contextual representations (Raza et al., 2017; Stefanelli et al., 2016) and play a role in spatial memory formation (Morales et al., 2021). However, neither activating nor inactivating somatostatin-positive interneurons before retrieval affected strategy choices, leaving spatial preference undisturbed in the dual solution paradigm. We then tested parvalbumin-positive interneurons, which represent mostly basket and chandelier cells located close to the granule cell layer that mediate inhibition near the soma and proximal dendrites of granule cells (Bartholome et al., 2020). Their chemogenetic activation in the DG has been shown to reduce anxiety and to enhance contextual fear memory extinction (Zou et al., 2016). In our experiments, activating PV-positive interneurons resulted in about 70% of mice choosing a cued response strategy. However, only 50% of the control vector-injected mice of the PV-Cre mouse line showed a preference for the spatial solution, suggesting a genetic background effect and different basal behavior of this mouse line in the dual solution paradigm. While it is intriguing to speculate that increased dDG inhibition by activating PV-positive interneurons may contribute to shifts in strategy choices, future experiments using different interventional approaches are necessary to answer this question.

Finally, in addition to manipulating specific interneuron populations, we used a chemogenetic vector that allowed for the inhibition of excitatory cells of the dDG (under control of the CamKIIa promotor). The use of an activating chemogenetic construct in these mice however had to be abandoned after pilot experiments, since the locomotor activity of the animals was low and side effects such as seizures were frequently observed. Focusing on inactivating DG excitatory cells before retrieval, we observed a shift towards a cue-preference. By contrast, inactivating the excitatory dDG cells before training did not induce such a shift (see Suppl. Fig. 5). Together, these findings suggest that either increasing (via GAD65 knock down and potentially also by blocking PV interneurons) or decreasing the excitability of the DG (via chemogentic inhibition of excitatory cells) both induces a shift towards cue-based strategies specifically in the retrieval phase of the dual solution paradigm. This suggests that an optimum excitation/ inhibition balance in the dDG is required for its proper functioning within a strategy choice network that may comprise the striatum and frontal cortical areas. Moreover, a recent study demonstrates that mossy cells of the hilus respond as well to objects explored in specific spatial configurations (GoodSmith et al., 2022). Mossy cells are the second excitatory cell population of the dDG next to granule cells. They are located in the hilus, receive inputs from granule cells as well as from back projections of CA3 and they target other mossy cells, granule but also basket cells. Depending on neuromodulatory inputs (i. e. acetylcholinergic inputs), mossy cells can provide a feed forward inhibition of granule cells, but also directly activate sparse granule cell populations (Schafman, 2016). They may therefore provide a switch for DG activity during the dual solution task depending on arousal and comprise an interesting target to study local circuits of the DG in future studies.

## 5. Conclusions and future perspectives

Setting out to investigate neuronal circuits underlying memory-based foraging strategies, we identified an unexpected role of the DG in mediating strategy selection during memory retrieval. Alterations in GAD65-mediated inhibition in the DG seem to provide a molecular mechanism of such an adaptive response. It is well known that the dDG is embedded in a functional network of decision-making processes, comprising for example the mPFC, the ACC and the anterior thalamic nucleus (Méndez-Couz et al., 2015), that may support its role as a hub for strategy selection. In fact, cFos-based regional activation analysis indicated altered functional interaction of DG and ACC in GAD65-/- mice during the dual solution task. Direct hippocampal inputs to the frontal cortex stem mainly from the ventral hippocampal CA1 areas but not the DG (Eichenbaum, 2017), but a knock down of GAD65 in the DG resulted in no alterations of excitability of CA1 or CA3 (see Suppl. Fig. 3 and 4). One promising relay station to be considered is the supramammillary nucleus (SUM), which projects bidirectionally to the ACC, to the PFC and specifically to the dorsal DG (Pan and McNaughton, 2004) and synchronizes its activity with the dDG during spatial memory retrieval (Li et al., 2020). Deconstructing this network and the contribution of such PFC-dDG-relay stations to spatial and cue-based information processing and retrieval will be an attractive task for future studies and may provide novel insights in networks of decision-making processes.

## Abbreviations

ACC: anterior cingulate cortex
BLA: basolateral amygdala
CA: Cornu ammonis
CeA: central amygdala
CNO: clozapine-N-oxide
CORT: corticosterone
dDG: dorsal dentate gyrus
DREADD: designer receptors exclusively activated by designer drugs
fEPSP: field postsynaptic potentials
GAD65: glutamic acid decarboxylating enzyme, 65 kDa isoform
GR: glucocorticoid receptors
LA: lateral amygdala
mPFC: medial prefrontal cortex
MR: mineralocorticoid receptors
PL/IL: Pre-/ Infralimbic cortex

## Acknowledgement

We thank Franziska Blitz, Antje Koffi von Hoff, Romina Wolter and Annika Lenuweit for excellent technical assistance and Gina M. Krause for assistance with the behavior experiments.

