## Supplementary Material for "Choosing memory retrieval strategies: a critical role for inhibition in the dentate gyrus"

### Supplementary Figures

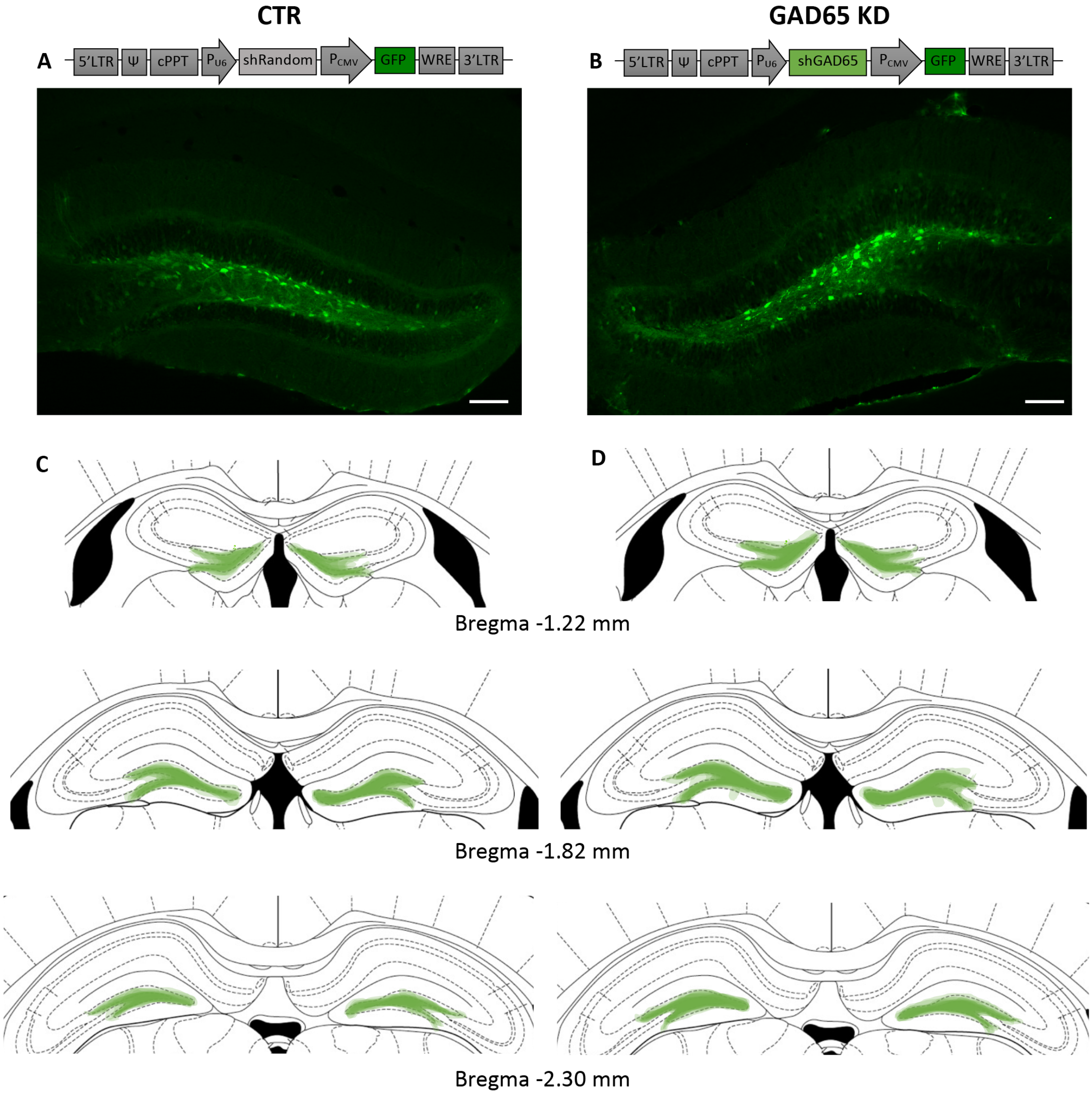


**Suppl. Fig. 1: GFP signal in GAD65 knock down experiment.** Mice were injected **(A)** with either a viral vector for expressing a random control shRNA sequence (CTR) or **(B)** a shRNA for knock down of GAD65 (GAD65 KD) into the dorsal dentate gyrus. Example microscopic photographs of the DG injected with CTR and GAD65 knock down vector are shown below the scheme of the viral vector design (scale bar: 100 µm). **(C+D)** GFP expression in each animal of the two groups was sketched at 3 levels of Bregma for the dorsal DG and overlays for each of the individual animals were created to illustrate spread of viral vector expression (in green).

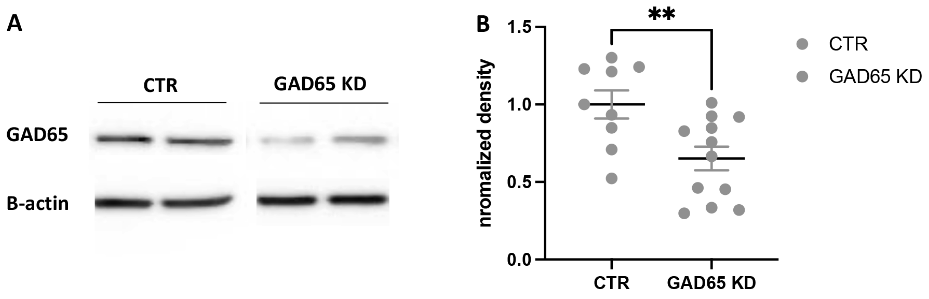


**Suppl. Fig. 2: GAD65 knock down efficiency in the mouse hippocampus.**
In 3 independent experiments, mice were injected in the dorsal dentate gyrus and ventral dentate gyrus/ CA3 region of the hippocampus (sh random control vector: CTR, n=9; sh GAD65 vector: GAD65 KD, n=12). 2 weeks later, 400 µm horizontal hippocampal slices were prepared and snap frozen in liquid nitrogen and stored until further preparation. For protein lysis frozen tissue samples were homogenized in 50 µl 1x SDS Buffer (62.5 mM tris-HCl, 2% SDS, 0.005% Bromophenolblue, 5% β-mercaptoethanol, 10% glycerol, all Sigma Aldrich, Germany) with freshly added proteinase inhibitor and phosphatase inhibitor cocktail (PhosStop, Roche Diagnostics, Germany). 10 µg of each sample was loaded on a 10% SDS-polyacrylamide gel for electrophoresis (SDS-PAGE). After semi-dry transfer (nitrocellulose membrane) and blocking in 5% milk with 0.1% Tween, membranes were incubated with primary antibodies over night at 4°C, using mouse α GAD65 1:1000 (ab26113, Abcam) and mouse anti β-actin 1:40000 (#A1978, Sigma Aldrich, Germany) as endogenous control. After incubation with complementary secondary antibody (anti ms IgG POD 1:10000, BM Chemilumenescence Western Blotting kit, Roche Diagnostics, Mannheim, Germany)), chemiluminescence was detected using Luminescence substrate (BM Chemiluminescence Western blot kit, Roche Diagnostics). Example blots are shown under **(A)**. The density of signals was analyzed with the Quantity One 1-D Analysis software (BioRad, Germany). Ratios between the optical density of target protein and the control protein β-actin were calculated for each sample and normalized to the mean density of the control group of each area. **(B)** Normalized mean densities of CTR and GAD65 KD slices were compared with t-tests after confirming normal distribution using a Shapiro-Wilk test. Across hippocampal subregions, GAD65 knock down reduced relative GAD65 protein expression levels to 65.2±0.07% on average compared to CTR injected tissue (normalized to 1; $T(19)=2.956$, $p=0.0081$), comparable to the efficiency yielded in the rat dorsal DG with the same construct (Tripathi et al., 2021).
All values mean ± sem. ** significant difference between groups, $p<0.01$.

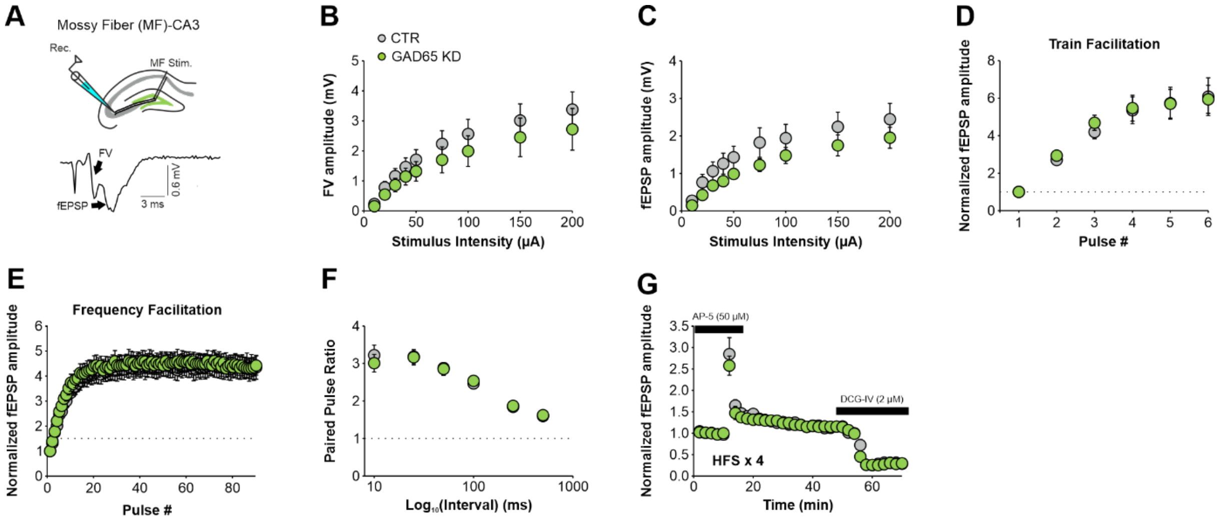


**Suppl. Fig. 3: GAD65 knock down in the dorsal dentate gyrus does not alter hippocampal mossy fiber-to-CA3 neurotransmission and plasticity.**

**(A)** Hippocampal scheme showing the stimulation and recording configuration for mossy fiber-to-CA3 synapse electrophysiology. Below is a representative trace illustrating presynaptic fiber volley (FV) and field excitatory postsynaptic potential (fEPSP). The recording electrode was placed at the stratum lucidum (SL) of CA3 and stimulation electrode was placed at the dentate hilus close the granule cell layer to stimulate mossy fibers (MF) as previously described (Madencioglu et al., 2021). After confirming stable excitatory fPSP (fEPSP) responses, input-output (I-O) curve was obtained which was followed by paired pulse, train facilitation (TF; 6 pulses at 20 Hz, repeated 3x at 0.05 Hz) and frequency facilitation (FF; 90 pulses at 1 Hz) protocols using the ~30-40 % of the maximum fEPSP amplitude. The amplitude of presynaptic fiber volley (FV) was calculated using the peak-to-peak amplitude (mV) of descending phase of the FV. For calculating the paired-pulse ratio, the magnitude of the second evoked potential was divided by the first one. To obtain TF and FF curves, the fEPSP amplitude values were normalized to the amplitude of the 1$^{st}$ fEPSP. To verify the recording of well-isolated MF-CA3 fEPSP responses, facilitation during the TF and FF protocols was analyzed. Slices were accepted for further analysis only when the last fEPSP amplitude was at least 250% of the 1st fEPSP amplitude. Analysis of I-O curves revealed no effect of GAD65 KD on the **(B)** presynaptic FV (CTR: n=11 slices; GAD65 KD n=11 slices) and **(C)** fEPSP amplitudes (CTR: n=11 slices; GAD65 KD: n=18 slices), thus indicating no change in baseline synaptic transmission. No effect on short term plasticity was evident when **(D)** train facilitation (CTR: n=9 slices; GAD65 KD n=11 slices), **(E)** frequency facilitation (CTR: n=7 slices; GAD65 KD n=10 slices) and **(F)** paired pulse ratios (CTR: n=11 slices; GAD65 KD n=11 slices) were compared between control (CTR) slices and GAD65 KD slices.

For high frequency stimulation (HFS)-induced long-term potentiation (LTP) first baseline recording was obtained for 10 min (Interval: 0.033 Hz; Pulse duration: 100 µs), then four HFS trains (100 pulses at 100 Hz with 20 s inter-train-intervals) were delivered. During baseline period and HFS delivery, slices were perfused with NMDA-R blocker AP-5 (50 µM) to isolate MF-induced NMDA-R independent LTP and prevent a possible contamination from association/commissural (A/C) synapses. Test pulses were delivered for consecutive 40 min to assess HFS-induced increase in fEPSP amplitudes. At the end of experiment, slices were washed with the group II metabotropic glutamate receptor agonist (2S, 2’R, 3’R)-2-(2’, 3’dicarboxycyclopropyl) glycine (DCG-IV, 2 µM, Tocris) for additional 30 min to confirm MF origin of the fEPSP responses. Data were only accepted when the reduction in fEPSP amplitude

was equal or more than 70%. The amplitude of fEPSP was analyzed via calculating the peak-to-peak amplitude (mV) of descending phase of the fEPSP. To measure the LTP induced by HFS, the amplitude values were normalized to the fEPSP responses obtained during the baseline period. **(G)** LTP remained unaltered in the MF-CA3 synapse (CTR: n=5 slices; GAD65 KD n=5 slices).
All data are reported as mean ± sem. Statistical comparison (B-G) was performed using two-way Repeated Measures ANOVA with Greenhouse-Geißer corrections for issues of sphericity when necessary (see the supplementary table 1 below for statistical information).

| **Supplementary Table 2 (Suppl. Fig. 4)** | **Main GAD65 KD effect** | **Stimulus Intensity, Interval or Pulse number** | **Interaction (GAD65 KD x Intensity) (GAD65 KD x Interval) (GAD65 KD x Pulse number)** |
|---|---|---|---|
| **(B) I-O curve (FV)** | F(1,20)=0.637, p=0.434 | F(1.074, 21.48)=41.30, p<0.0001 | F(8,160)=0.412, p=0.913 |
| **(C) I-O curve (fEPSP)** | F(1,20)=1.655, p=0.213 | F(1.393, 27.87)=65.85, p<0.0001 | F(8,160)=0.676, p=0.712 |
| **(D) Train Facilitation** | F(1,18)=0.027, p=0.872 | F(1.111, 19.99)=52.38, p<0.0001 | F(5,90)=0.174, p=0.972 |
| **(E) Frequency facilitation** | F(1,15)=0.063, p=0.805 | F(2.095, 31.43)=65.32, p<0.0001 | F(89,1335)=0.429, p>0.999 |
| **(F) Paired Pulse Ratios** | F(1,20)=0.017, p=0.898 | F(2.524, 50.48)=68.96, p<0.0001 | F(5,100)=0.393, p=0.852 |
| **(G) Long-term potentiation** | F(1,8)=0.687, p=0.431 | F(2.176, 17.41)=100.4, p<0.0001 | F(34,272)=0.598, p=0.964 |

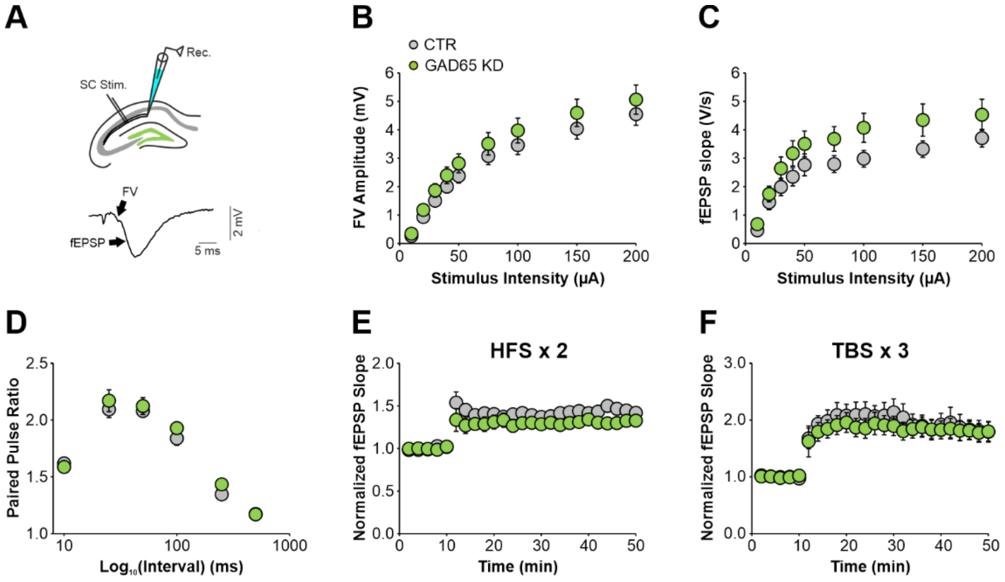


**Suppl. Fig. 4: GAD65 knock down in the dorsal dentate gyrus does not alter hippocampal Schaffer collateral-to-CA1 neurotransmission and plasticity.**

**(A)** Hippocampal scheme showing the stimulation and recording configuration for Schaffer collateral (SC)-to-CA1 synapse electrophysiology. Below is a representative trace illustrating presynaptic fiber volley (FV) and field excitatory postsynaptic potential (fEPSP). The recording electrode was placed at the dendritic stratum radiatum (SR) of CA1 and stimulation electrode was placed at the proximal SR of CA1 to stimulate Schaffer collaterals (SC) as previously described (Annamneedi et al., 2021, 2018). After confirming stable fEPSP responses, an input-output (I-O) curve was obtained which was followed by a paired pulse protocol using the ~50 % of the maximum fEPSP amplitude analogous to DG and CA3 electrophysiology. The slope of negative-going fEPSP was analyzed by calculating the slope (V/s) between the 20 and 80% of the fEPSP amplitudes. The amplitude of presynaptic fiber volley (FV) was calculated using the peak-to-peak amplitude (mV) of descending phase of the FV. For calculating the paired-pulse ratio, the magnitude of the second evoked potential was divided by the first one.

Analysis of input-output (I-O) curves revealed no effect of GAD65 KD on the **(B)** presynaptic FV (CTR: n=24 slices; GAD65 KD n=21 slices) and **(C)** fEPSP amplitudes (CTR: n=24 slices; GAD65 KD n=21 slices) indicating no change in baseline synaptic transmission. No effect on short term plasticity was evident when **(D)** paired pulse ratios were compared between control slices and GAD65 KD slices (CTR: n=24 slices; GAD65 KD n=21 slices).

LTP was induced either with two HFS (2 trains of 100 pulses at 100 Hz with 20 s inter-train-interval) or three theta burst stimulations (each TBS including 10 trains of 10 pulses at 100 Hz with 200 ms inter-train-interval and 10 s inter-TBS-interval). Similar to CA3 electrophysiology baseline recording (Interval: 0.033 Hz; Pulse duration: 100 µs) was obtained for 10 min and test pulses were delivered for 40 min after HFS or TBS protocols to measure LTP. To measure the LTP induced by HFS or TBS, the slope values were normalized to the fEPSP responses obtained during the baseline period. Both **€** high frequency stimulation (HFS)-induced (CTR: n=9 slices; GAD65 KD n=8 slices) and **(F)** theta burst stimulation (TBS)-induced long-term potentiation (LTP) (CTR: n=7 slices; GAD65 KD n=8 slices) remained unaltered in the SC-CA1 synapse.

All data are reported as mean ± sem. Statistical comparison (B-F) was performed using two-way Repeated Measures ANOVA with Greenhouse-Geißer corrections for issues of sphericity when necessary (see the supplementary table 2 below for statistical information).

| **Supplementary Table 2 (Suppl. Fig. 4)** | **Main GAD65 KD effect** | **Stimulus Intensity, Interval or Pulse number** | **Interaction (GAD65 KD x Intensity) (GAD65 KD x Interval) (GAD65 KD x Pulse number)** |
|---|---|---|---|
| **(B) I-O curve (FV)** | F(1,43)=1.033, p=0.315 | F(1.131, 48.62)=208.7, p<0.0001 | F(8,344)=0.520, p=0.841 |
| **(C) I-O curve (fEPSP)** | F(1,43)=2.667, p=0.101 | F(2.372, 102.0)=65.87, p<0.0001 | F(8,344)=1.152, p=0.328 |
| **(D) Paired Pulse Ratios** | F(1,43)=0.566, p=0.456 | F(3.420, 147.1)=166.0, p<0.0001 | F(5,215)=0.715, p=0.613 |
| **(E) Long-term potentiation (HFS)** | F(1,15)=2.529, p=0.133 | F(3.319, 49.79)=28.88, p<0.0001 | F(24,360)=1.300, p=0.159 |
| **(F) Long-term potentiation (TBS)** | F(1,13)=0.017, p=0.207 | F(2.560, 33.28)=29.76, p<0.0001 | F(24,312)=0.406, p=0.995 |

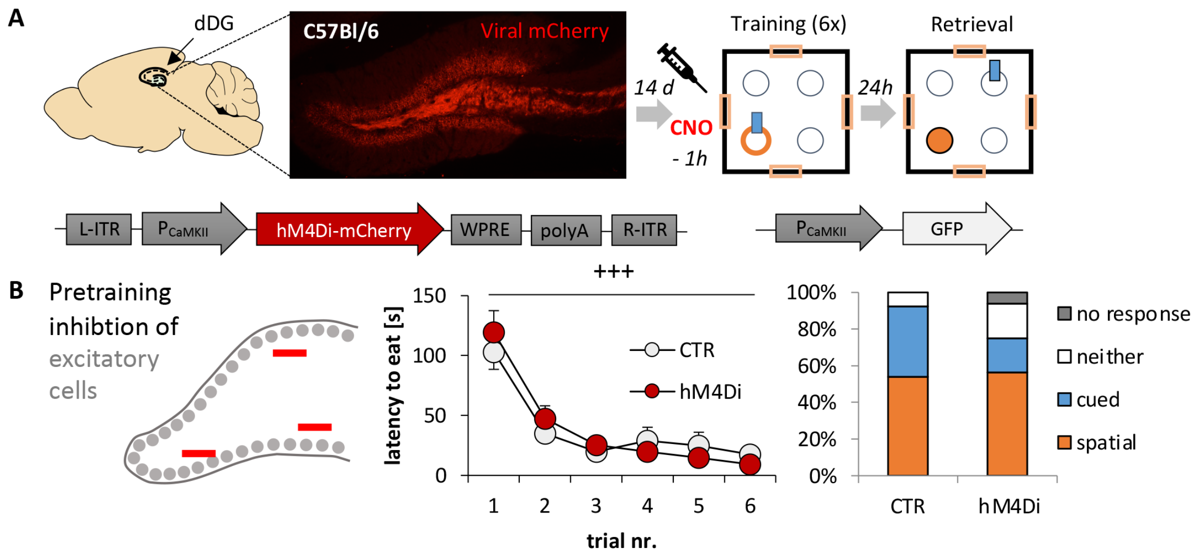


**Suppl. Fig. 5: No effect of pre-training chemogenetic inhibition of excitatory cells in the DG.** (A) Scheme of chemogenetic inhibition of excitatory DG cells via DREADDs. (B) Inhibiting DG activity before training (CamKII-hM4Di, n=13; CamKII-CTR, n=16), had no effect on learning the reward location (Repeated Measure ANOVA with Greenhouse-Geißer correction for trial $F(2.049,51.226)=37.225$, $p<0.001$; trial x group interaction $F(2.049,51.226)=0.723$, $p=0.494$; group $F(1,25)=0.128$, $p=0.724$). Moreover, both groups maintained a spatial preference during retrieval 24h later, when excitability in the dDG projections were normal again ($X^2(2)=2.466$, $p=0.481$). All values mean ± sem. +++ significant learning effect over trials, $p<0.001$.

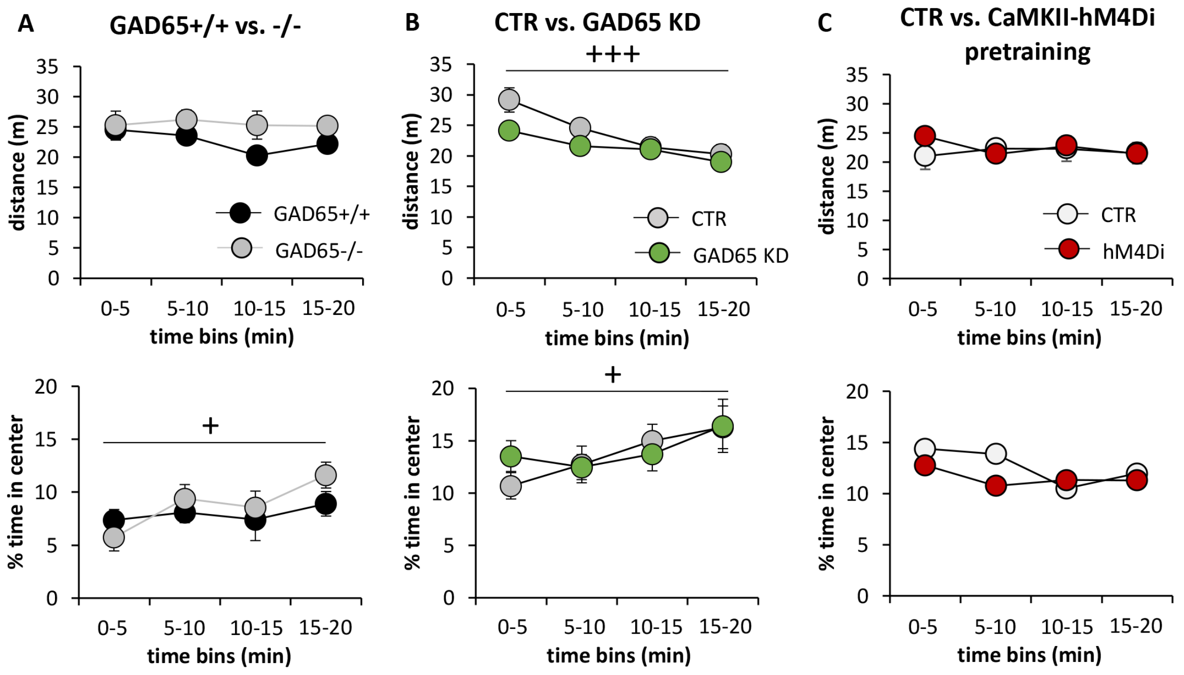


**Suppl. Fig. 6: Activity and anxiety in the OF before dual solution training.**
Around one hour before the dual solution training took place in an open field arena, all animals were habituated to the open field for 20 min. The open field data can be used to assess potential differences in activity and anxiety-like behavior via measuring the total distance the animals covered and the % time spent in the center, respectively. **(A)** Neither in GAD65 deficient animals (GAD65+/+ n=9; GAD65-/- n=7) nor **(B)** after local knock down of GAD65 in the dorsal dentate gyrus (CTR: n=16; GAD65 KD n=18) or **(C)** by activation of excitatory dorsal dentate gyrus cells via chemogenetic inhibition under the CaMKII-Promotor (CTR n=16, hM4Di n=13) were any changes induced in the open field behavior.
All values mean ± sem. Statistical comparison was performed using two-way repeated measures ANOVA for 5 min time bins with Greenhouse-Geißer corrections for issues of sphericity when necessary (see the supplementary table 3 below for statistical information). + significant effect for time, p < 0.05+++ p<0.001.

| **Supplementary Table 3 (Suppl. Fig. 6)** | **group** | **time** | **time x group** |
|---|---|---|---|
| **(A) GAD65+/+ vs. GAD65-/-** ***distance*** | F(1,13)=3.622, p=0.79 | F(1.441, 18.730)=0.751, p=0.444 | F(1.441, 18.730)=0.667, p=0.477 |
| **(A) GAD65+/+ vs. GAD65-/-** ***% time in center*** | F(1,13)=0.348, p=0.565 | F(1.789, 23.262)=5.155, p=0.017 | F(1.789, 23.262)=1.375, p=0.271 |
| **(B) CTR vs. GAD65 KD** ***distance*** | F(1,32)=2.229, p=0.145 | F(1.746, 56.463)=19.837, p<0.001 | F (1.746, 56.463)=2.267, p=0.119 |
| **(B) CTR vs. GAD65 KD** ***% time in center*** | F(1,32)=0.049, p=0.826 | F(3,96)=3.299, p=0.024 | F(3,96)=0.685, p=0.563, p=0.271 |
| **(C) CTR vs. hM4Di** ***distance*** | F(1,27)=0.193, p=0.664 | F(2.021,54.57)=0.4, p=0.753 | F(2.021,54.57)=1.037, p=0.381 |
| **(C) CTR vs. hM4Di** ***% time in center*** | F(1,27)=0.223, p=0.64 | F (3,81)=1.261, p=0.293 | F(3,96)=0.672, p=0.572 |
